# On the unfounded enthusiasm for soft selective sweeps

**DOI:** 10.1101/009563

**Authors:** Jeffrey D. Jensen

## Abstract

Underlying any understanding of the mode, tempo, and relative importance of the adaptive process in the evolution of natural populations is the notion of whether adaptation is mutation-limited. Two very different population genetic models have recently been proposed in which the rate of adaptation is not strongly limited by the rate at which newly arising beneficial mutations enter the population. This review discusses the theoretical underpinnings and requirements of these models, as well as the experimental insights on the parameters of relevance. Importantly, empirical and experimental evidence to date challenges the recent
enthusiasm for invoking these models to explain observed patterns of variation in humans and *Drosophila*.

## INTRODUCTION

Identifying the action of positive selection from genomic patterns of variation has remained as a central focus in population genetics. This owes both to the importance of specific applications in fields ranging from ecology to medicine, but also to the desire to address more general evolutionary questions concerning the mode and tempo of adaptation. In this vein, the notion of a soft selective sweep has grown in popularity in the recent literature, and with this increasing usage the definition of the term itself has grown increasingly vague. A soft sweep does not reference a particular population genetic model per se, but rather a set of very different models that may result in similar genomic patterns of variation. Further, it is a term commonly used in juxtaposition with the notion of a hard selective sweep, the classic model in which a single novel beneficial mutation arises in a population and rises in frequency quickly to fixation. Patterns expected under the hard sweep model have been well described in the literature (see reviews of [1,2]; and Figure 1), and consist of a reduction in variation surrounding the beneficial mutation owing to the fixation of the single haplotype carrying the beneficial, with resulting well-described skews in the frequency spectrum (e.g., [3-5]) and in patterns of linkage disequilibrium (e.g., [6-8]). Indeed, part of the recent popularity of soft sweeps comes from the seeming rarity of these expected hard sweep patterns in many natural populations (see for example [9-11]).

**Figure 1:**
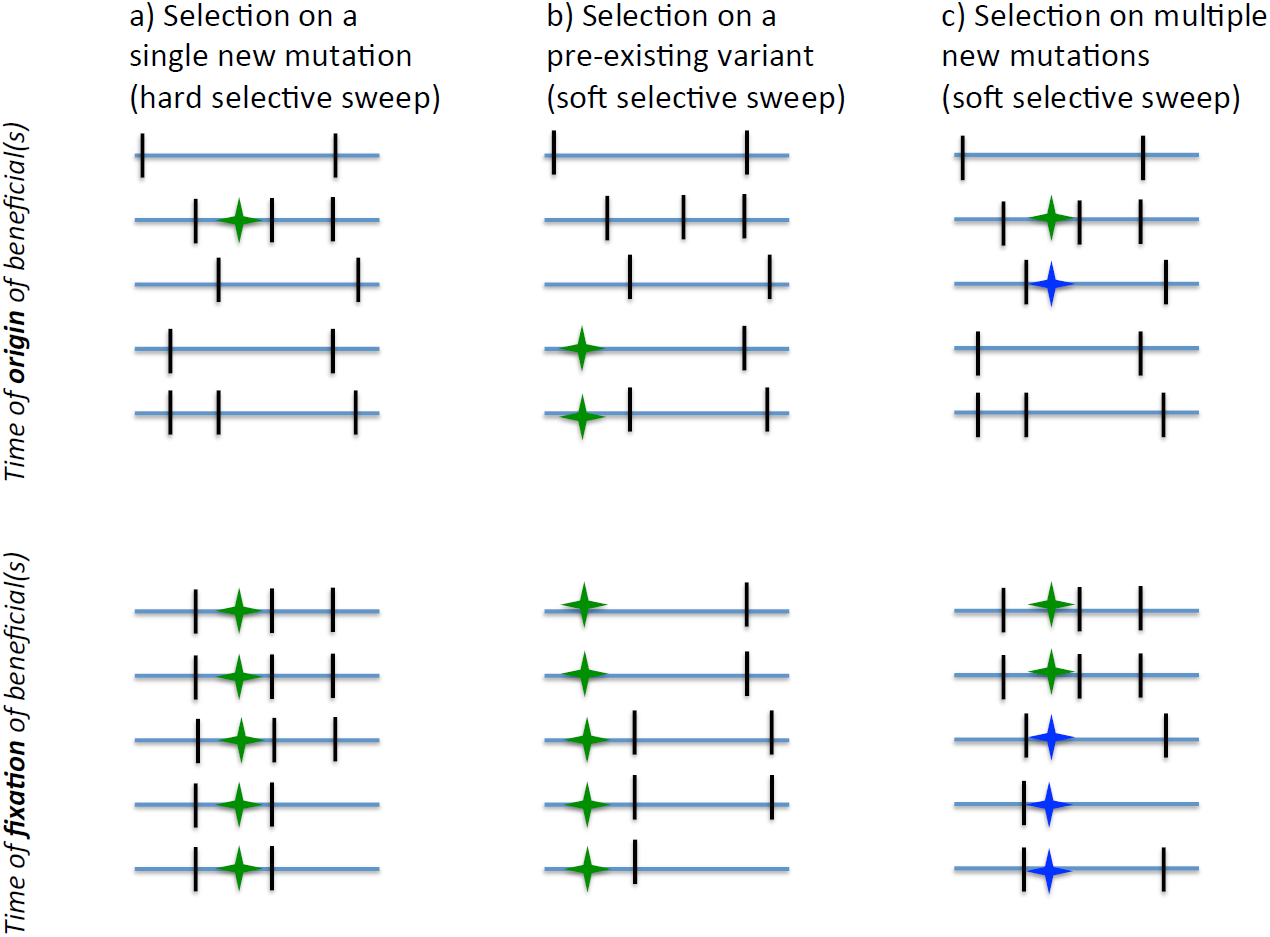
Cartoon representation of three population genetic models of selection Each column represents a model: (a) classic hard selective sweep (i.e., selection on a newly arising beneficial mutation), (b) selection on standing variation (i.e., a pre-existing variant which has become beneficial), and (c) selection on multiple new mutations (i.e., a second beneficial mutation arises prior to the fixation of the first). The first row represents the time of the origin of the beneficial(s), in which five sampled chromosomes are shown with blue lines, each of which carries neutral mutations (black dashes) and some of which carry the beneficial mutation (given by the green star, where the blue star represents a second independently arising beneficial in the ‘multiple new mutation’ model). The second row represents the time of fixation of the beneficial mutation. In the hard sweep model (a), the beneficial mutation as well as closely linked neutral variation has been brought to fixation, while more distant sites may only be brought to intermediate frequency owing to recombination. In the pre-existing variant model (b), the beneficial mutation may carry the haplotypes on which it was segregating prior to the shift in selective pressure each to some intermediate frequency. In the multiple new mutation model (c), each independent beneficial mutation may carry to intermediate frequency the haplotype on which it arose. Thus, models b) and c) produce a qualitatively similar end result.

In terms of patterns of variation, the primary difference between soft and hard selective sweeps lies in the expected number of different haplotypes carrying the beneficial mutation or mutations, and thus in the expected number of haplotypes that hitchhike to appreciable frequency during the selective sweep, and which remain in the population at the time of fixation. This key difference results in different expectations in both the site frequency spectrum and in linkage disequilibrium, and thus in the many test statistics based on these patterns (see Figure 1). Owing to this ambiguous definition, a number of models have been associated with producing a soft sweep pattern – including selection acting on previously segregating mutations, and multiple beneficials arising via mutation in quick succession (see review of [11], and Box 1).

##### BOX 1: Overview of two soft selective sweep models

The first of these models associated with soft selective sweeps, popularized by Hermisson & Pennings [13], is one in which a given beneficial mutation previously segregated in the population neutrally (or at an appreciable frequency under mutation-selection balance), and thus existed on multiple haplotypes at the time of the selective shift in which the mutation became beneficial. In this way, a single beneficial mutation may carry multiple haplotypes to intermediate frequency, while itself becoming fixed (see Figure 1). Though Hermisson & Pennings associated this model with the term ‘soft sweep’, the model of selection on standing variation has been long considered in the literature. Orr & Betancourt [15] considered the model in some detail as will be discussed below, as did Innan & Kim [89]. Indeed, Haldane [90] also discussed the possibility of selection acting on previously deleterious mutants segregating in the population.

A second commonly invoked model associated with soft selective sweeps, also popularized by Pennings & Hermisson [31], is one in which multiple beneficial mutations independently arise in short succession of one another – such that a second copy arises via mutation prior to the selective fixation of the first copy. While this model was first envisioned as consisting of multiple identical beneficial mutations (i.e., the identical change occurring at the same site), it has since been considerably expanded to include any mutation which produces an identical selective effect (e.g., if all loss-of-function mutations produce an equivalent selective advantage, a large number of possible mutations may be considered as being identically beneficial [91]). The similarity to the standing variation model described above, and thus their shared association with the notion of a soft sweep, is simply that these multiple beneficial mutations may arise on different haplotypes, and thus also sweep different genetic backgrounds to intermediate frequency (see Figure 1).

However, apart from shared expected patterns of variation, these two population genetic models are very different. Selection on standing variation requires that the beneficial mutation segregate at appreciable frequency in the pre-selection environment, whereas the multiple beneficial model requires a high mutation rate to the beneficial genotype. One important point which will be returned to throughout is the distinction between the relevance of these models themselves, and the likelihood of these models resulting in a hard (i.e., single haplotype) versus a soft (i.e., multiple haplotype) selective sweep at the time of fixation. Below I will discuss what is known from theory regarding these models, and what is known from experimental evolution and empirical population genetic studies regarding the values of the key parameters dictating their relevance. I conclude by arguing that the recent enthusiasm for invoking soft sweeps to explain observed patterns of variation is likely to be largely unfounded in many cases.

## SELECTION ON STANDING VARIATION

### Expectations and assumptions

A good deal is known from the theory literature regarding the likelihood of selection on standing variation, and the parameter space of relevance for this model. Kimura’s [12] diffusion approximation gave the fixation probability of an allele (*A*) segregating in the population at frequency (*x*) with selective advantage (*s*_*b*_):

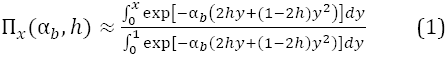

where *h* is the dominance coefficient and α_*b*_ = 2*N*_*e*_*s*_*b*_ (where *N*_*e*_ is the effective population size). Following Hermisson and Pennings [13], if selection on the heterozygote is sufficiently strong (i.e., 2h*α*_*b*_ ≫ (1-2h) /2h), this may be approximated as:

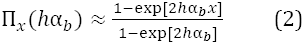

where for a new mutation (i.e., *x* = 1/2*N*), we find Haldane’s [14] result that the fixation probability is approximately twice the heterozygote advantage 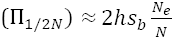.

Comparing this result with the situation in which the beneficial allele was previously segregating neutrally (i.e., *x* > 1/2*N*), Hermisson and Pennings [13] obtained the following approximation:

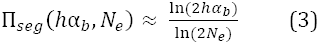

demonstrating that the fixation probability is much greater for beneficials that already begin at an appreciable frequency (because of their lower probability of being lost by drift) – though this condition may be misleading, as the segregation of a neutral allele at intermediate frequency is indeed already an unlikely event (which could be considered by integrating over the distribution of neutral variant frequencies). They further approximate the probability that the population adapts from standing variation (sgv) as:

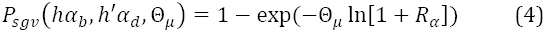

where *R*_*α*_ = 2*hα*_*b*_/(2*h*′α_*d*_ + 1) measures the selective advantage of the allele in the new relative to the old environment (where α_*b*_ is the selective effect in the new environment and α_*d*_ in the previous environment, and *h* and *h’* are the dominance coefficients in the new and previous environments respectively), and Θ_*μ*_ is the population mutation rate. Thus, for an allele at mutation-selection balance (i.e., *x* = Θ_*μ*_/2*h*′α_*d*_), this probability is simply ≈ 1 – exp(–Θ_*μ*_*h*α_*b*_/*h*′α_*d*_).

Thus, the key parameters for understanding the likelihood of a model of selection on standing variation involves knowing the pre-selection fitness of beneficial mutations (i.e., prior to becoming beneficial), and relatedly the pre-selection frequency of beneficial mutations (again, prior to becoming beneficial). Below, I will briefly review what is known from both experimental and empirical studies regarding these parameters in the handful of instances in which we have good inference.

### Empirical evidence regarding pre-selection allele frequency

Orr and Betancourt [15] previously considered this model and reached a similar result as Hermisson and Pennings [13] – namely, a soft sweep from standing variation only becomes feasible when the mutation has a non-zero probability of segregating at an appreciable frequency at the time of the selective shift (i.e., the beneficial mutation was previously neutral or slightly deleterious and segregating under drift at relatively high frequency, it was maintained at appreciable frequency by balancing selection prior to the selective shift, etc…). Indeed, they provide a direct calculation for the probability that multiple copies of the beneficial allele (*X*) exist, conditional on fixation of the beneficial mutation:

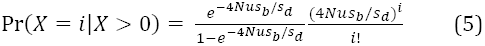

where *s*_*d*_ is the selection coefficient prior to the shift, *s*_*b*_ is the selection coefficient after the shift, and *Nu* is the population mutation rate.

With this, Orr and Betancourt made the notable observation that even if selection is acting on standing variation instead of a new mutation, a single copy is nonetheless surprisingly likely to sweep to fixation (i.e., producing a hard, rather than a soft, sweep from standing variation). Indeed, they demonstrate that multiple copy fixations become more likely than single copy fixations from standing variation only when 4*Nus*_*b*_/*s*_*d*_> 1. For reasonable parameter estimates, they calculate that the allele must be present in many copies in the population before obtaining an appreciable probability of sweeping multiple copies of the beneficial mutation to fixation. For example, from Orr and Betancourt, for *N* = 1x10^4^, *s*_*d*_ = 0.05, *s*_*b*_ = 0.01, *u* = 10^-5^, and *h* = 0.2, 96% of the time a single copy will fix in the population, despite 20 copies segregating at mutation-selection balance prior to the shift in selection pressure. For these parameters the population size must be in excess of *N* = 1.5x10^5^ (thus more than 300 copies segregating at mutation-selection balance) before multiple copies are more likely upon fixation than a single copy.

Revisiting this model, Przeworski et al. [16] more explicitly examined the frequency at which a mutation must be segregating prior to the shift in selection pressure, before multiple haplotypes would likely be involved in the selective sweep. They found that a hard sweep is likely when *x* < 1/2*N*_*e*_*s*_*b*_ (consistent with the simulated exampled from Orr and Betancourt above). Thus, taking the mutation-selection balance frequency given above, we may conclude that a hard sweep (i.e., involving a single haplotype) is likely from standing variation when Θ_*μ*_/2*h*′α_*d*_ < 1/2*N*_*e*_*s*_*b*_ (see Figure 2). An important distinction is again necessary here. While the parameter requirement mentioned above concerns the likelihood of a soft sweep from standing variation, it further suggests that we are unlikely to have statistical resolution when attempting to distinguish between a hard sweep on a new mutation vs. a hard sweep on a rare previously standing variant.

**Figure 2:**
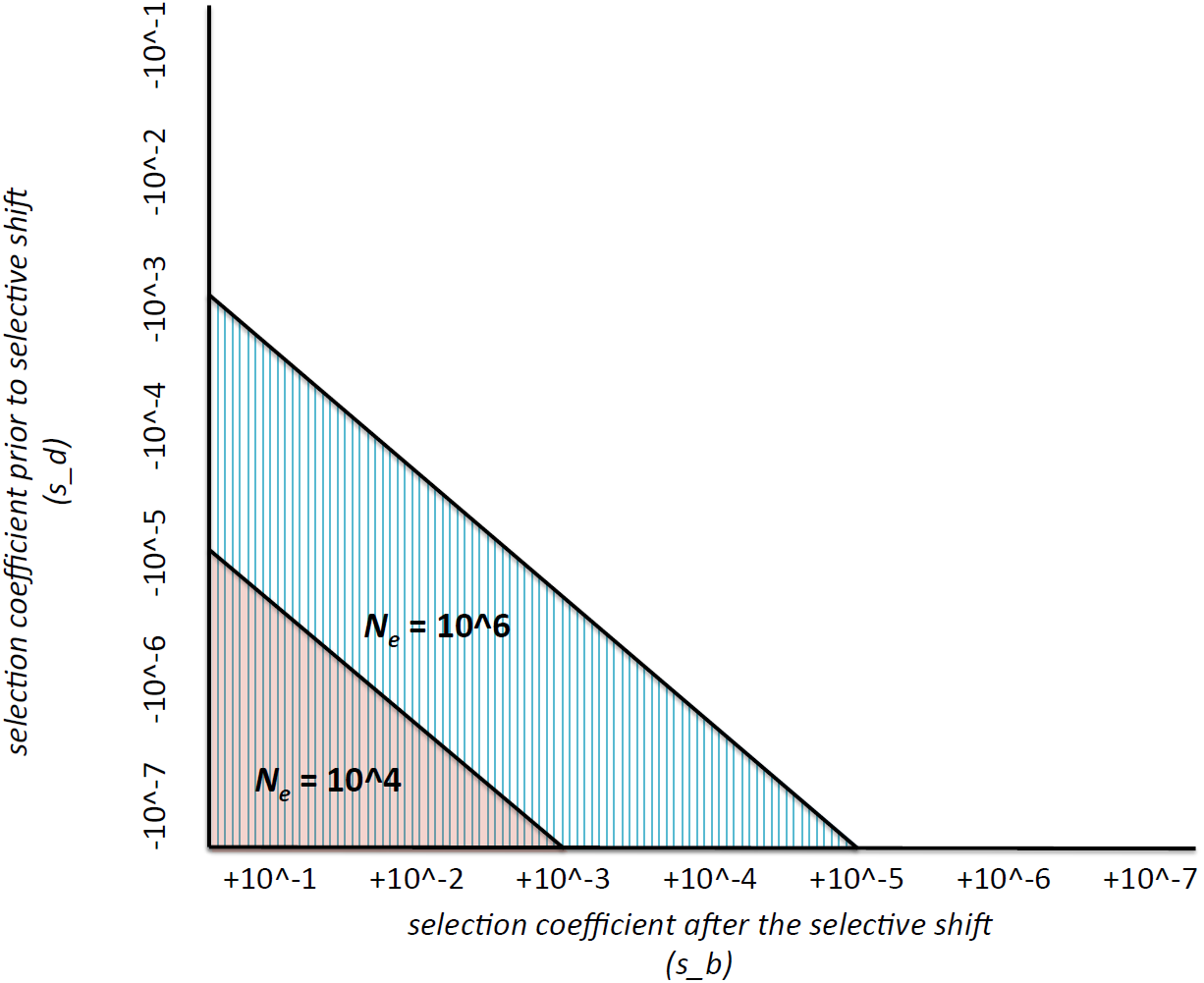
The conditions under which a soft sweep from standing variation becomes possible The Y-axis represents the selection coefficient prior to the selective shift (i.e., given by negative selection coefficients) and the X-axis is the selection coefficient after the shift in selective pressure (i.e., given by positive selection coefficients). The area under each line represents the parameter space for which such a soft sweep is feasible for two different effective population sizes – one human-like (104, given by pink shading) and one *Drosophila*-like (10^6^, given by vertical blue lines). As shown, the effect prior to the selective shift must be nearly neutral or weakly deleterious in order for the allele to segregate at an appreciable frequency, and the effect post selective shift must be strongly beneficial. As described in the text, this inference rests on the argument that Θ_*μ*_/2*h*′α_*d*_ must be greater than 1/2*N*_*e*_*s*_*b*_ for a soft selective sweep from standing variation to become likely, where here Θ_*μ*_ = 10^-8^ and *h*′ = 1.

As an empirical example of the above point, one of the most widely cited and discussed examples of selection on standing variation surrounds the *Eda* locus in Sticklebacks [17]. With evidence for selection reducing armor plating in freshwater populations compared to the ancestral heavily plated marine populations, the authors sequenced marine individuals in order to estimate the allele frequency of the freshwater adaptive low plate morphs, with estimates ranging from 0.2%-3.8%. While the low plate morph is likely deleterious in marine populations (potentially suggesting that it is at mutation-selection balance), migration from the marine environment may indeed serve as an important source of variation for local freshwater hard sweeps. However, as noted by the authors, it is difficult to separate this hypothesis from that of local freshwater adaptation on new mutations, followed by back migration of locally adapted alleles in to the marine population. Similarly, related arguments have been made for a large amount of rare standing variation being responsible for the quick and persistent response of phenotypic traits to selection in the quantitative genetics literature (for a helpful review, see [18]). However, as with the above example, distinguishing rare standing ancestral variation from newly accumulating mutations has also been a topic of note (e.g., [19]). Regardless of these caveats, hard sweeps from rare standing variants segregating at mutation-selection balance in ancestral populations, rather than on *de novo* mutations alone, appear to be an important and viable model of adaptation.

### Experimental and empirical evidence regarding the cost of beneficial mutations

Based on the simple and enlightening result of Orr and Betancourt, it is reasonable to ask, for cases in which we have reasonably strong functional evidence of adaptation, what we know about the value of 4*Nus*_*b*_/*s*_*d*_, as this will dictate the likelihood of a hard vs. a soft sweep from standing variation. There are two fields from which we may obtain insight – experimental-evolution studies in which the selective effects of mutations may be precisely measured under controlled environmental conditions, and empirical population genetic studies in which inference can be drawn about the selective effect of functionally validated mutations in the presence and absence of a given selective pressure.

First, there is a rich literature in experimental evolution from which we can draw. In a recent evaluation of the distribution of fitness effects in both the presence and absence of antibiotic in the bacterium *Pseudomonas fluorescens*, Kassen and Bataillon [20] found that of the 665 resistance mutations isolated, greater than 95% were deleterious in the absence of antibiotic treatment. In populations of yeast raised in both standard and challenging environments (in this case, high temperature and high salinity), Hietpas et al. [21] identified a handful of beneficial mutations in each of the challenge environments, all of which were deleterious under standard conditions, with some even being lethal in the absence of the selective pressure. Foll et al. [22] in investigating the evolution of oseltamivir resistance mutations in the influenza A virus, identified 11 candidate resistance mutations, with the one functionally validated mutation (H274Y) having been demonstrated to be deleterious in the absence of drug pressure (see also [23-24]).

Second, there are a small but increasing number of examples from natural populations where we have both a functionally validated beneficial mutation for which we understand the genotype-phenotype connection, as well as inference on the selection coefficient both in the presence and absence of a given selective pressure. One such example is the evolution of cryptic coloration in wild populations of deer mice [25]. In the Nebraska Sand Hills population, population genetic and functional evidence has been found for positive selection acting on a small number of mutations modifying different aspects of the cryptic phenotype, all contained within the *Agouti* gene region [26]. Three lines of population genetic evidence suggest that selection began acting on these mutations when they arose (i.e., selection on a de novo mutation): 1) the beneficial mutations appear to be carried on single haplotypes (though, as discussed above, selection on standing variation may indeed often only result in a single haplotype fixation), 2) the beneficial mutations have not been sampled off of the Sand Hills region (i.e., the mutation is unlikely to have been segregating at appreciable frequency in the ancestral population prior to the formation of the Sand Hills), and 3) using an approximate Bayesian approach, the age of the selected mutation has been inferred to be younger than the geological age of the Sand Hills [25]. In addition, ecological information pertaining to this phenotype exists as well. Performing a predation experiment with clay models, Linnen et al. [26] demonstrated a strong selective advantage of crypsis – with conspicuous models being subject to avian predation significantly more than cryptic models. This result suggests that if the beneficial phenotype currently present in the Sand Hills was indeed present in the ancestral population, it was likely to be strongly deleterious.

Other notable examples exist in the empirical literature as well. For example, Agren et al. [27] mapped quantitative trait loci (QTL) for 398 recombinant inbred lines of Arabidopsis derived from crossing locally adapted lines from Sweden and Italy. Their results suggests a small number of locally adaptive genomic regions, and that in many cases the locally adaptive change was deleterious in the alternate environment. Performing a meta-analysis on a wide range of antibiotic resistance mutations in pathogenic microbial populations, Melnyk et al. [28] found that across eight species and fifteen drug treatments, resistance mutations were widely found to be deleterious in the absence of treatment (i.e., in 19/21 examined studies). At the *Ace* locus of *D. melanogaster*, four described variants conferring varying degrees of pesticide resistance have been described, all of which are strongly deleterious in the absence of this pressure (with deleterious selection coefficients ranging from -5% to -20%; see [29]).

Thus, given the required pre-selection frequency necessary to result in a soft rather than a hard sweep, it is fair to say that this combination of results provide poor support for the relevance of soft sweeps from standing variation in the populations examined. However, it is worth noting that such studies likely represent an ascertainment bias towards traits that strongly affect the phenotype, thus making them amenable for ecological and laboratory study. Assuming a relationship between the observed phenotypic and underlying selective effects, it may well be that beneficial mutations of small effect (which are more difficult to study and thus under-represented in the literature) may be those more likely to have only weakly deleterious effects in the absence of a given selective pressure.

## MULTIPLE COMPETING BENEFICIALS

### Expectations and assumptions

The second model that I will discuss is that of multiple competing beneficial mutations of identical selective effects. As opposed to the standing variation model described above in which there was a single mutational origin of the beneficial mutation, this model posits multiple beneficial mutations of independent origin arising in quick succession of one another. Importantly, these independent beneficial mutations must necessarily arise on different haplotypes. If the second independently arising identical beneficial mutation arises on the same background as the first, the resulting pattern would simply be that of a hard sweep as they would be impossible to distinguish (i.e., a single haplotype would be swept).

Thus, the expected number of haplotypes and their frequency distributions is a necessary consideration. This expectation is given by Ewens’ sampling formula (see [30]), for the case of no recombination. Given a mutation rate of Θ_*b*_ to the *B* allele, the probability to find *k* haplotypes occurring *n*_*1*_, *n*_*2*_, …*n*_*k*_ times in a sample of size *n* is given as:

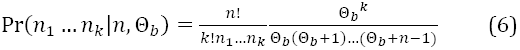

For *k* = 1 and *n*_*1*_ = 1, Pennings and Hermisson [31] thus wrote the upper bound for the probability of such a soft sweep as:

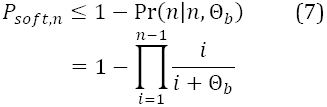

Hence, the probability of a soft sweep from multiple competing beneficials is naturally dependent on the beneficial population mutation rate. As they describe, the beneficial mutation rate must be extremely large before this model becomes likely (with Θ_*b*_ > 0.04 before even two haplotypes would be swept with an appreciable probability, for *α* = 10,000). In other words, within the relatively fast sojourn time of a beneficial mutation (*T*_*fix*_ ≈ 4*N*_*e*_log(*α*)/*α*), multiple identical beneficial mutations must arise on separate haplotypes, escape drift (where, once again, the probability of fixation of each new independent mutation is given as 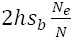), and ultimately fix in multiple copies.

Thus, the key parameter for understanding the likelihood of a model of competing beneficials involves knowing the mutation rate to the beneficial genotype. This latter clarification gives rise to an additional parameter – namely, the size of the mutational target available for creating an identical beneficial mutation. If only a single mutational change is possible, the target size is one base pair. If, as given as an example in the Introduction, any loss-of-function mutation within a coding region produces an identical selective effect, the target size may be dozens of base pairs or more. Below, I will briefly review what is known from both experimental and empirical studies regarding these parameters in the handful of instances in which we have good inference.

### Experimental and empirical evidence regarding the beneficial mutation rate

To understand the beneficial mutation rate is fundamentally to understand the distribution of fitness effects – that is, the proportions of newly arising mutations that are beneficial, neutral, and deleterious. Characterizing this distribution has spawned a long and rich literature among both theoreticians and experimentalists. Fisher [32] had already considered the probability that a random mutation of a given phenotypic size would be beneficial, concluding that adaptations must consist primarily of small effect mutations. Kimura [33] recognized one difficulty with this conclusion, noting that while small effect mutations may be more likely to be adaptive, large effect mutations have a higher probability of fixation. Thus, Kimura argued that in fact intermediate effect mutations may be most common in the adaptive process. Orr [34] gained an important additional insight - given any distribution of mutational effects, the distribution of factors fixed during an adaptive walk (i.e., the sequential accumulation of beneficial mutations) is roughly exponential. An important by-product of this result is the notion that the first step of an adaptive walk may indeed be quite large (in agreement with Fisher’s Geometric Model).

Efforts to quantify the shape of the distribution of fitness effects (DFE) and characterize the beneficial mutation rate have come largely from the experimental evolution literature. One common feature amongst this work is the use of extreme value theory (EVT) (see review of [35]). Because the DFE of new mutations is generally considered to be bimodal (e.g., [36]) – consisting of a strongly deleterious mode and a nearly-neutral mode, beneficial mutations represent the extreme tail of the mode centered around neutrality. One particular type of extreme value distribution – the Gumbel type (which contains a number of common distributions including normal, lognormal, gamma, and exponential) – has been of particular focus beginning with Gillespie [37].

Recently, experimental efforts have begun to better characterize the shape of the true underlying distribution in lab populations experiencing adaptive challenges (see review of [38]). Though the fraction of beneficial mutations relative to the total mutation rate is indeed small, providing good support for the assumptions of EVT, the exact shape of the beneficial distribution varies by study. Sanjuan et al. [39] found support for a gamma distribution using site-directed mutagenesis in vesicular stomatitis virus. Kassen and Bataillon [20] found support for an exponential distribution assessing antibiotic resistance mutations in *Pseudomonas*. Rokyta et al. [40] found support not for the Gumbel domain but rather for a distribution with a right-truncated tail (i.e., suggesting that there is an upper bound on potential fitness effects), using two viral populations. MacLean and Buckling [41], again using *Pseudomonas*, argued that an exponential distribution well-explained the data when the population was near optimum, but not when the population was far from optimum, owing to a long tail of strongly beneficial mutations. Schoustra et al. [42], working in the fungus Aspergillus, demonstrated that adaptive walks tend to be short, and characterized by an ever-decreasing number of available beneficial mutations with each mutational step taken. One important caveat in such experiments however is that they commonly begin from homogenous populations. Thus, while providing a good deal of insight in to the underlying DFE, they are far from direct assessments of the relative role of single *de novo* beneficial mutations in adaptation.

Whole genome time-sampled sequencing is also shedding light on the fraction of adaptive mutations. Examining resistance mutations in the influenza virus both in the presence and absence of oseltamivir (a common drug treatment), Foll et al. [22] identified the single and previously described resistance mutation (i.e., H274Y [43]) as well as 10 additional putatively beneficial mutations based upon duplicated experiments and population sequencing, suggesting a fraction of 11/13588 potentially beneficial genomic sites in the presence of drug treatment, or 0.08% of the genome. But perhaps the most specific information currently available regarding beneficial mutation rates comes from experiments in which all mutations may be generated individually (as opposed to mutation-accumulation studies) and directly evaluated across different environmental conditions (see [21,44]). Within this framework, Bank et al. [45] recently evaluated all possible 560 individual mutations in a sub-region of a yeast heat shock protein (Hsp90) across 6 different environmental conditions (standard, as well as temperature and salinity variants), identifying few beneficials in the standard environment, and multiple beneficials associated with high salinity. In order to quantify this shift, the authors fit a Generalized Pareto Distribution (GPD), using the shape parameter (*κ*) to summarize the changing DFE – with the Weibull domain fitting the less-challenging environments (i.e., demonstrating that the DFE is right-bounded, suggesting that the populations are near optimum), and the Frechet domain fitting the challenging environment (i.e., a heavy-tailed distribution owing to the presence of strongly beneficial mutations, potentially suggesting that the population is more distant from optimum).

Thus, despite some small but important differences between these conclusions, there is general support for a model in which newly arising mutations take a bimodal distribution, with the extreme right tail of this distribution representing putatively beneficial mutations. In other words, the beneficial mutation rate is likely a very small fraction of the total mutation rate. Given the requirements of the multiple competing beneficials model (i.e.,Θ_*b*_ > 0.04), this would seemingly make the model only of relevance to populations of extremely large N_*e*_*μ*, as in perhaps certain viral populations (see [46]). Indeed, in an attempt to argue for the relevance of these models in *Drosophila*, Karasov *et al*. [29] claim an effective population size in *Drosophila* that is orders of magnitude larger than commonly believed (i.e., *N*_*e*_ > 10^8), despite the great majority of empirical evidence to the contrary (see review of [47]).

### Experimental and empirical evidence regarding the mutational target size

However, despite the above conclusion, if the mutational target size is not a single site, but rather a large collection of sites, this value may become more attainable for a wider array of species. As with the above section, the most abundant and reliably validated information comes from experimental evolution. While still imperfect as these studies do not necessarily reflect all potential available beneficial solutions (i.e., mutation accumulation studies can only draw inference on the mutations which happen to occur during the course of the experiment, and studies using direct-mutagenesis have thus far only evaluated sub-genomic regions). Returning to the examples given in the section above, we may ask what are the functional requirements of the identified beneficial mutations. In studying adaptation to the antibiotic rifampicin in the pathogen *Pseudomonas*, MacLean and Buckling [41] demonstrated that the beneficial mutations identified are consistent with known molecular interactions between rifampicin and RNA polymerase – as the antibiotic binds to a small pocket of the β-subunit of RNAP, in which only 12 amino acid residues are involved in direct interaction. Wong et al. [48] investigated the genetics of adaptation to cystic fibrosis-like conditions in Psuedomonas both in the presence and absence of fluoroquinolone antibiotics, describing a small number of stereotypical resistance mutations in DNA gyrase. Examining the evolution of oseltamivir resistance in the influenza A virus, Foll et al. [22] described a similar story, in which the small handful of putatively beneficial resistance mutations are concentrated in HA and NA, with the single characterized resistance mutation being shown to alter the hydrophobic pocket of the NA active site, thus reducing affinity for drug.

Again considering the natural populations for which we have solid genotype-phenotype information and about which we understand something about the nature of adaptation acting on these mutations, let us consider a few examples. Describing wide-spread parallel evolution on armor plating in wild threespine sticklebacks, Colosimo et al. [17] demonstrated the Ectodysplasin (EDA) signaling pathways to be repeatedly targeted for modifications to this phenotype with a high degree of site-specific parallel evolution. Looking across 14 insect species that feed on cardenolide-producing plants, Zhen et al. [49] also noted repeated bouts of parallel evolution for dealing with this toxicity not only confined to the same alpha subunit of the sodium pump (ATPα), but in the great majority of cases to the same two amino acid positions. Examining adaptation to pesticide resistance in *Drosophila*, four specific point mutations in the *Ace* gene have been identified which result in resistance to organophosphates and carbonates (see [29]). Cryptic coloration has also been a fruitful area, with specific mutations in the *Mc1r* and *Aguoti* gene regions having been described as the underlying cause of adaptation for crypsis in mice of the Arizona / New Mexico lava flows [50], Nebraska Sand Hills [25] and the Atlantic coast [51], as well as in organisms ranging from the Siberian mammoth [52] to multiple species of lizards on the White Sands of New Mexico [53-54] (and see review of [55] for further examples).

Thus, for the handful of convincing genotype-to-phenotype examples in the literature, the adaptive mutational target size appears small, a result which would appear to be biologically quite reasonable.

### The effects of selection on linked sites

However, even if the mutational target size is sufficiently large such that a model of competing beneficials becomes feasible, it becomes necessary to consider interference between these segregating selected sites. It is helpful to consider three relevant areas of the parameter space: a) beneficial mutations of identical selective effects arising in a low recombination rate region, potentially allowing for a soft sweep; b) beneficial mutations of differing selective effects arising in a low recombination rate region, where the most strongly beneficial likely outcompetes the others producing a hard sweep; or c) multiple beneficial mutations occurring in a high recombination rate region, allowing for a hard sweep of the most beneficial haplotype (i.e., the recombinant carrying the most beneficial mutations).

Hill and Robertson [56] explicitly considered the probability of fixation for two segregating beneficial mutations. Confirming the arguments of Fisher [32], they demonstrated that selection at one locus indeed interferes with selection at the alternate locus, reducing the probability of fixation at both sites – with the conclusion being that simultaneous selection at more than one site reduces the overall efficacy of selection (see also [57-58]).

This effect is clearly a function of the amount of recombination between the selected sites. If the sites are independent there is no such effect, and if they are tightly linked the effect will be very strong (Figure 3). While it is difficult to generalize this information, for the current empirical data available discussed above, it appears both likely and biologically reasonable to consider that mutations conferring identical selective effects may indeed be occurring within a narrow genomic region (e.g., mutations within the drug binding pocket HA in influenza virus in response to drug, within RNAP in *Pseudomonas* in response to antibiotic, at the *Agouti*/*Mc1r* locus in vertebrates for color modifications, at the Ace locus of *Drosophila* for pesticide resistance, at the *Eda* locus in Sticklebacks for armor modifications, etc…).

**Figure 3:**
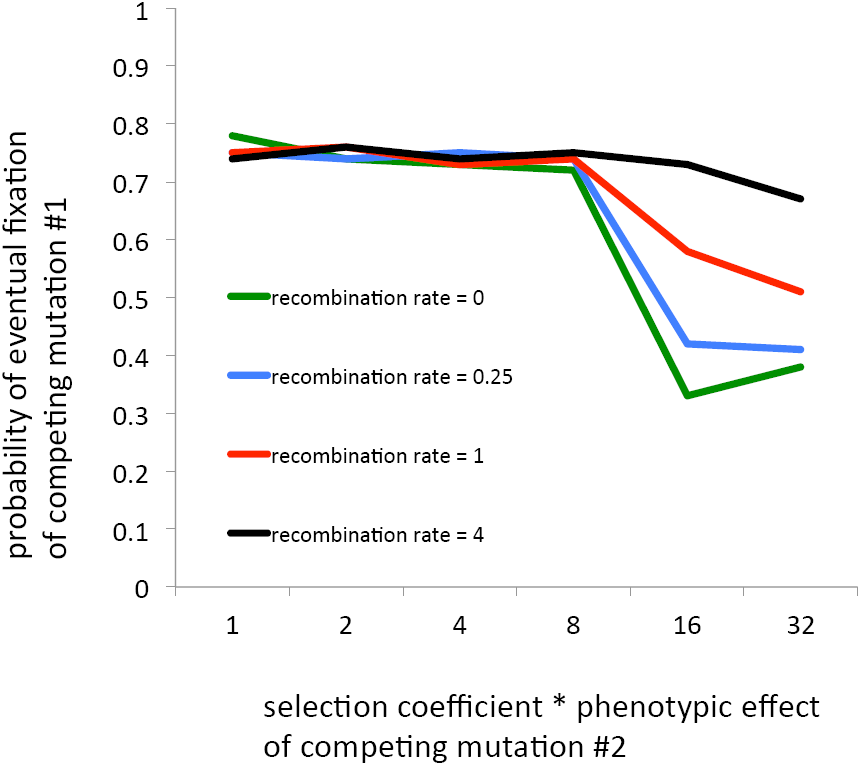
Probability of fixation of a first beneficial mutation under a scenario of two competing beneficial mutations Representation of the results of Hill and Robertson [56] (modified from their Figure 2). On the Y-axis is the probability of fixation of the first beneficial mutation, and on the X-axis is the selection coefficient multiplied by the phenotypic effect of the second (and thus competing) beneficial mutation. Here, the selective effect of mutation #1 is 8, on the scale given on the X-axis, and each beneficial mutation is assumed to be at a 10% frequency in the population. The four lines represent four different recombination rates between the two sites. As shown, if the effect of the second site is weaker than the first (i.e., < 8), the first beneficial retains a high probability of fixation and the second site will be lost unless it can recombine on to a common haplotype. However, as the effect of the second beneficial mutation becomes stronger, the probability of fixation of the first beneficial mutation decreases rapidly – a phenomenon that is increasingly strong with tighter genetic linkage between the sites. Thus, for rare to moderate recombination, and a strong beneficial effect of the second site, the probability of fixation of the first beneficial mutation is halved even for this simple case of only two competing beneficials.

Examining the extent of this effect by simulation, Comeron et al. [59] found that the effect becomes stronger as 1) the sites become more weakly beneficial, 2) the recombination rate is decreased, and 3) the number of selected sites increases (consistent with the results of [60-61]). However, as long as there is linkage between the sites, the probability of fixation decreases rapidly as the number of selected sites grows, even for very strong selection. Examining the effect of two competing beneficial mutations in the presence of recombination analytically, Yu and Etheridge [62] further demonstrate the relative likelihood of an ultimate single haplotype fixation.

Thus, while a large mutational target size may in principle increase the relevance of this model, it results in a scenario still requiring a large beneficial mutation rate, necessitates that these beneficial mutations escape initial stochastic loss, and finally, owing to interference, results in a decreased probability of fixation for each competing beneficial relative to independence. Again invoking results from experimental evolution, in a highly informative recent study by Lee and Marx [63], the authors demonstrate the strong effects of clonal interference in replicated populations of *Methylobacterium extorquens* – identifying as many as 17 simultaneous beneficial mutations existing in a population which may rise in frequency initially, only to be lost owing to competition with an alternate and ultimately successful single beneficial mutation, in what they termed repeated ‘failed soft sweeps.’

As a natural population example, Hedrick [64] discusses the multiple identified malaria resistance variants identified in humans, and makes the case that in the continued presence of malaria, single variants are highly likely to ultimately fix at the cost of losing other competing and currently segregating beneficial resistance mutations, owing to measured selection differentials. Similarly, at the previously discussed *Ace* locus of *D. melanogaster*, a single of the four identified resistance mutations was found to confer 75% resistance to pesticide, two mutations confer 80% resistance, and three mutations confer full resistance – again suggesting that a single haplotype carrying multiple beneficial mutations will likely ultimately result in a hard selective sweep. Both of these observations, along with the results of Lee and Marx [63], suggest that multiple competing beneficial mutations may indeed be a likely model, but a soft sweep from multiple beneficials is unlikely owing to non-equivalent selective effects between the mutations (or the haplotypes carrying these mutations). Thus, as with the model of selection on standing variation above – the model of competing beneficial mutations itself has good empirical and experimental support for being relevant, but a hard sweep rather than a soft sweep appears as the more likely outcome given our current understanding of the parameters of relevance.

## PERSPECTIVE & FUTURE DIRECTIONS

Apart from the considerations discussed in the sections above, and conditional on the unsubstantiated assumption that adaptive fixations are common, the absence of hard sweep patterns in many natural populations has led some to conclude that soft sweeps must be the primary mechanism of adaptation, with a recent popularity for invoking these models in the human and *Drosophila* literature. However, as argued above, this assumption is poorly supported, and theoretical and experimental insights to date suggest that soft sweeps from standing variation or from multiple beneficial mutations for populations of this size are unlikely. This argument itself is of course somewhat circular, as quantifying the fraction of adaptively fixed mutations, and the proportion of newly arising beneficial mutations, is indeed one of the central focal points of population genetics, and is far from resolved as discussed. Thus, assuming a very large fraction of adaptive fixations in order to quantify the fraction of adaptive fixations is rather self-defeating.

A quite separate point has also been neglected in this literature. Namely, the power of existing tests of hard selective sweeps to identify these patterns within demographically complex populations (a category that certainly includes humans and *Drosophila*). Biswas and Akey [65] examined the consistency between methods used for conducting genomic scans for beneficial mutations in humans. Results differed dramatically, ranging from 1799 genes identified by Wang et al. [66] to 27 genes identified by Altshuler et al. [67]. Perhaps even more striking, of the six studies examined, there was virtually no overlap in the genes identified. For example, of the 1799 genes identified by Wang et al. [66], 125 overlap with the scan of Voight et al. [68], 47 from Carlson et al. [69], 5 from Altshuler et al. [66], 4 from Nielsen et al. [70], and 40 from Bustamante et al. [71]. Additionally, the recent review by Crisci et al. [2], summarizing estimates of the rate of adaptive fixation in *Drosophila*, noted that the inferred genomic rate differs by two orders of magnitude between studies (from *λ* = 1.0E-12 [72] to *λ* = 1.0E-11 [73-74] to *λ* = 1.0E-10 [75] - where 2*N*_*e*_*λ* is the rate of beneficial fixation per base pair per 2*N*_*e*_ generations).

Evaluating the performance of these statistics has thus remained an important question, and over the past decade numerous researchers have demonstrated low power under a wide-range of neutral non-equilibrium models (e.g., [76-80]). More recently, Crisci et al. [81] specifically evaluated the ability of the most widely used and sophisticated tests of selection via simulation (Sweepfinder [70], SweeD [80], and OmegaPlus [82]), to identify both complete and incomplete hard selective sweeps under a variety of demographic models of relevance for human and *Drosophila* populations. The results are troubling, with the true positive rate rarely exceeding 50% even under equilibrium models, and being considerably worse for models of moderate and severe population size reductions (Figure 4). Furthermore, the false positive rate was often in excess of power, particularly for models of population bottlenecks. Though not conclusive, this indeed suggests a troubling potential interpretation for the lack of overlap between the above mentioned genomic scan studies.

**Figure 4:**
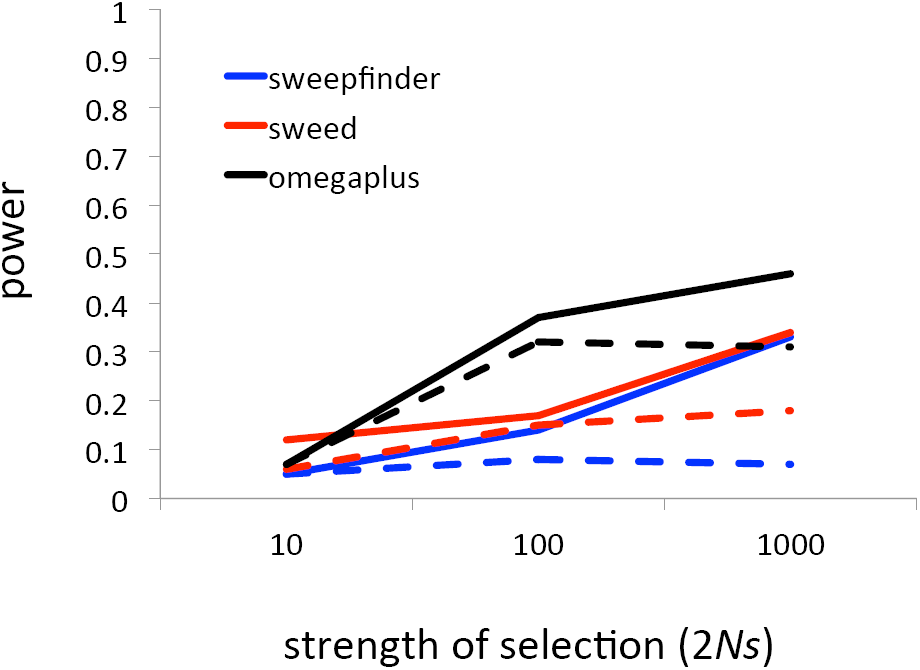
Statistical power to detect hard selective sweeps Representation of the results of Crisci et al. [81] examining the power of three of the most commonly-used approaches for detecting hard selective sweeps – Sweepfinder (blue), Sweed (red), and Omegaplus (black). On the Y-axis is the power of the test statistic, and on the X-axis is the strength of selection. Two models are plotted: 1) a selective sweep in an equilibrium population (given by the solid lines), and 2) a selective sweep in a moderately bottlenecked population, in which the population is reduced to 10% of its former size 0.2 4*N* generations in the past (given by the dashed lines). In all cases, *N*_*e*_ = 10^4^ and the selective event occurred 0.01 4*N* generations in the past. As shown, even in equilibrium populations the power scarcely reaches 50% even for strong selection, while in mildly bottlenecked populations the power approaches nominal levels for frequency spectrum based statistics (i.e., Sweed and Sweepfinder) and drops to 30% for the better performing LD-based approaches (i.e., Omegaplus).

If nothing else, these results demonstrate that absence of evidence is not evidence of absence for the hard sweep model –implying that we only have minimal power to detect even very recent and very strong hard selective sweeps in these populations, and essentially no power for the great majority of the parameter space. However concerning these results may be, it is important that the field has made the effort to quantify the performance of the test statistics designed for detecting hard sweeps – defining Type-I and Type-II error and examining performance in demographic models both with and without selection. This scrutiny has yet to be brought to soft sweep expectations and statistics. Before these models can be reasonably invoked as explanations for observations in natural populations, we need to similarly understand the ability of neutral demographic models to replicate soft sweep patterns, quantify our ability to identify soft sweeps from standing variation and from multiple beneficial mutations in non-equilibrium populations, and understand the effects of relaxing current assumptions involving linkage and epistasis (i.e., for selection on standing variation the assumption is made that a single beneficial mutation will have the same selective effects on all genetic backgrounds, and the multiple beneficial model assumes that there are no epistatic interactions between co-segregating mutations). Early efforts have been made in some of these areas, with recent work examining basic expectations of these models under fluctuating effective population sizes, resulting in a further description of how population size changes may result in the ultimate fixation of a single beneficial mutation [83].

In conclusion, the wide-array of genomic patterns of variation that may be accounted for by models associated with soft selective sweeps has allowed adaptive explanations to proliferate in the literature, and be invoked for a larger subset of genomic data. However appealing this may be, these models in fact carry with them very specific and well-understood parameter requirements. Further, the ability of alternate models to produce these patterns needs to be more carefully weighed in future studies, particularly given preliminary findings concerning similar patterns produced under both neutral demographic models [81] and models of background selection [84]. Indeed, alternative models of positive selection have also been suggested to produce qualitatively similar patterns – including hard selective sweeps in subdivided populations exchanging migrants (e.g., [85-87]), and polygenic adaptation (e.g., [88]).

Finally, while examples in the literature are accumulating in support of the models themselves (e.g., selection on standing variation at the *Eda* locus of Sticklebacks or selection on multiple beneficials at the *Ace* locus of *Drosophila*), there is very little evidence of soft sweep fixations, with the best empirical and experimental examples to date almost universally pointing to hard sweep fixations under these models. This appears to primarily be owing to the low pre-selection allele frequency of the standing variants (which are seemingly often deleterious prior to the shift in selective pressure), and to the selective differential between competing beneficial mutations (or between the haplotypes carrying the beneficial mutations) resulting in the ultimate fixation of only a single haplotype. Thus, while the models themselves certainly deserve further attention, theoretical, empirical, and experimental results to date suggest that the field ought to take greater caution when invoking soft selective sweep fixations, as hard sweep fixations (be it from models of selection on new mutations, standing variation, or competing beneficial mutations) seem to remain as the most likely outcome across a wide parameter space relevant for many current populations of interest.

## Acknowledgements

I would like to thank Chip Aquadro, Roman Arguello, Dan Bolon, Margarida Cardoso Moreira, Brian Charlesworth, Laurent Excoffier, Adam Eyre-Walker, Joanna Kelley, Tim Kowalik, Anna-Sapfo Malaspinas, Bret Payseur, Molly Przeworski, Nadia Singh, Wolfgang Stephan, and Alex Wong for helpful comments and suggestions on an earlier version. I would also like to thank the authors of Melnyk, Wong & Kassen for sharing their manuscript while in review. I would finally like to thank members of the Jensen Lab for insightful comment and discussion throughout the writing process, in particular Claudia Bank, Anna Ferrer Admetlla, Matthieu Foll, Stefan Laurent, Louise Ormond, Cornelia Pokalyuk, and Nick Renzette. JDJ is funded by grants from the Swiss National Science Foundation, and a European Research Council (ERC) Starting Grant.

### Competing financial interests

The author declares no competing financial interests.

